# Ancient biomolecules suggest learned foraging strategy in extinct cave bears

**DOI:** 10.1101/2024.09.23.614392

**Authors:** Ioana N. Meleg, Federica Alberti, Dorothée G. Drucker, Magdalena Năpăruş-Aljancic, Angelica Feurdean, Marius Robu, Marius Vlaicu, Yuichi I. Naito, Adina Boroneanț, Marin Cârciumaru, Elena C. Nițu, Michael Hofreiter, Hervé Bocherens, Axel Barlow

**Affiliations:** Emil G. Racoviță Institute, Babeș-Bolyai University, Clinicilor 5-7, 400006, Cluj-Napoca, Romania; “Emil Racoviță” Institute of Speleology, Romanian Academy, Calea 13 Septembrie, nr. 13, 050711, Sector 5, Bucharest, Romania; Department of Bioinformatics and Genetics, Swedish Museum of Natural History, Stockholm, Sweden; Centre for Palaeogenetics, Svante Arrhenius väg 20C, 10691 Stockholm, Sweden; Institute for Biochemistry and Biology, Faculty for Mathematics and Natural Sciences, University of Potsdam, Karl-Liebknecht-Str. 24-25, 14476, Potsdam, OT Golm, Germany; Senckenberg Centre for Human Evolution and Palaeoenvironment (S-HEP) at the University of Tübingen, Hölderlinstraße 12, 72074, Tübingen, Germany; Karst Research Institute ZRC SAZU, Titov trg 2, SI-6230 Postojna, Slovenia; Tular Institute, Oldhamska 8A, SI-4000, Kranj, Slovenia; Department of Physical Geography, Goethe University, Altenhöferallee 1, 60438 Frankfurt am Main, Germany; Research Institute of the University of Bucharest, The Earth, Environmental and Life Sciences Division, 1 B.P. Hașdeu St., 50567, Bucharest, Romania; Department of Geosciences, Biogeology, University of Tübingen, Hölderlinstraße 12, 72074, Tübingen, Germany; Central Research Institute of Electric Power Industry (CRIEPI), 1646 Abiko, Abiko-shi, Chiba 270-1194, Japan; ”Vasile Pârvan” Institute of Archaeology, Romanian Academy, 11 Henri Coandă St, Bucharest, Romania; “Princely Court” National Museum Târgovişte, Museum of Human Evolution and Technology in Palaeolithic, 7 Justiţiei Street, Târgovişte 130017, Dâmboviţa County, Romania; School of Environmental and Natural Sciences, Bangor University, Bangor, LL57 2DG UK

**Keywords:** palaeogenomes, behavioural ecology, cave bears, dietary variation, palaeoclimate, stable isotopes

## Abstract

Studying the behavioural ecology of long-extinct species is challenging due to the difficulty in measuring the behavioural phenotype and correlating this with genetic and environmental factors. However, a multidisciplinary approach integrating isotope analysis of diet and ancient DNA analysis of genetic relationships offers a potential framework to test the proximate causes of dietary preferences. Our study focuses on Late Pleistocene cave bears from the Romanian Carpathians. Stable isotope analysis of bone collagen reveals substantial lifetime variation in food plant preferences among individuals. We find that bears with similar diets do not cluster according to their population structure, sex, time period, climatic conditions, or location. This disconnect suggests that diet preference in cave bears is not genetically inherited, and instead that individuals adapted their diets based on foraging experience. This integrative approach opens new avenues for understanding Pleistocene animal behaviour, leveraging ancient biomolecules synergistically to reveal insights otherwise inaccessible.

## Background

For over a century, scientists have grappled with the question of whether certain behaviours are inherited or acquired through learning (Galton, 1895). Thorpe’s definition of learning encompasses the process in which an individual’s behaviour adapts and/or responds as a consequence of their experiences (Thorpe, 1963). Extensive research has documented learning in many extant vertebrates, particularly in the context of foraging behaviour and vocalisations (Hoppitt & Laland, 2008; Whiten, 2017). However, long-extinct animals present a unique challenge in this regard. Although insights into their behavioural ecology have been made possible by combining palaeontological and ancient DNA evidence (Allentoft et al., 2015; Fortes et al., 2016; Huynen et al., 2010; Pečnerová et al., 2017), the proximate causes of their behaviour, in general, remain unknown. To address this challenge, an appropriate approach for studying the roles of learning and inheritance in the behaviour of extinct animals would require quantifying both variation in behaviour and genetic structure from their fossil remains.

Known for the high abundance of their fossil remains in caves across Europe, cave bears are one of the most extensively studied extinct Pleistocene species (e.g. Gretzinger et al., 2019; Hofreiter et al., 2004; Knapp et al., 2009; Pacher & Stuart, 2009; Stiller et al., 2010). Cave bears were large bodied bears that were widespread through Eurasia during the Late Pleistocene, and went extinct around 25,000 years ago (Pacher & Stuart, 2009; Terlato et al., 2019). They form the sister group of the brown bear and polar bear clade, from which they diverged around 1.5 million years ago (Barlow et al., 2021). Their diet has been the focus of several palaeoecological studies using stable isotopes. These reveal a herbivorous diet that varies geographically and between populations, with some cave bears exhibiting bulk collagen *δ*^15^N values exceeding 7‰), similar to those of exclusive herbivores like woolly mammoths and certain horses (Bocherens, 2019; Krajcarz et al., 2016; Naito et al., 2020; Pérez-Ramos et al., 2020).

Comparatively few studies have investigated the relationship between cave bear diet and their population genetics. A notable study found evidence of dietary niche partitioning between two morphologically discernible mitochondrial clades that are also quite distinct on the nuclear level (Barlow et al., 2018, 2021) but occurred in sympatry in two neighbouring caves in Austria (Bocherens et al., 2011). However, for other mitochondrial clades, no evidence of niche partitioning has been found at their zones of contact (Münzel et al., 2011), although it has to be noted that in this case representatives of these mitochondrial clades were found to be phylogenetically similar at the nuclear level (Barlow et al., 2018, 2021).

The most dramatic isotopic differentiation among cave bears across their entire geographical range occurred in the Romanian Carpathians. Here, the isotope signatures of their bone collagen separate into two distinct categories based on nitrogen and carbon stable isotope ratios. One group of bears has low *δ*^15^N and *δ*^13^C values, which is typical of all cave bears in the rest of Europe, whereas the second has high *δ*^15^N and low *δ*^13^C values, a pattern unique to the Romanian Carpathians (Bocherens, 2019; Robu et al., 2013, 2018). The isotopic signature obtained from adult bone collagen integrates data across an extensive timeframe, capturing dietary trends over several years leading up to death in long-lived large animals (Münzel et al., 2014; Huja & Beck, 2008). Thus, it remains unaltered by variations in seasons or the specific timing of the individual’s demise, and instead reflects an averaged value throughout the adult’s lifespan (Bocherens, 2015). Amino acid (glutamate and phenylalanine) nitrogen isotope analysis recently revealed that this high *δ*^15^N isotopic signature in bulk collagen reflects niche partitioning on different food plants exhibiting variation in *δ*^15^N (Naito et al., 2020).

In this study, we utilise palaeogenome data to investigate the factors underlying these distinct dietary niches. Specifically, we test for dietary niche partitioning by geographical location, population affinity, age of the samples, and sex. If none of these factors is found to be associated with the patterns of dietary variation, we hypothesise that these bears instead acquired their foraging behaviours during their lifetimes based on individual experience, fitting the definition of a learned behaviour.

## Methods

### Samples

Cave bear bones were sampled across Romania, including from well-studied caves containing important cave bear deposits of the Western Romanian Carpathian Mountains (i.e Apuseni Mountains: Igrița, Ciur Izbuc, Ciur Ponor, Meziad, Ferice, Măgura, Coliboaia and Onceasa caves; Banat Mountains: Răsuflătoarei, 2 Mai, Șelitrari and Climente I caves) and of the Southern Romanian Carpathians (i.e. Cioclovina Uscată, Cornetul Satului, Stogu, Cioarei de la Boroșteni, Muierilor and Colțul Surpat caves). We additionally sampled previously unstudied cave bear deposits in the Eastern Romanian Carpathians (i.e. Cave number 7 from Lelici, hereafter Lelici). Samples from Adam Cave, Dobrogea were also analysed (Figure 1).

**Figure 1.**
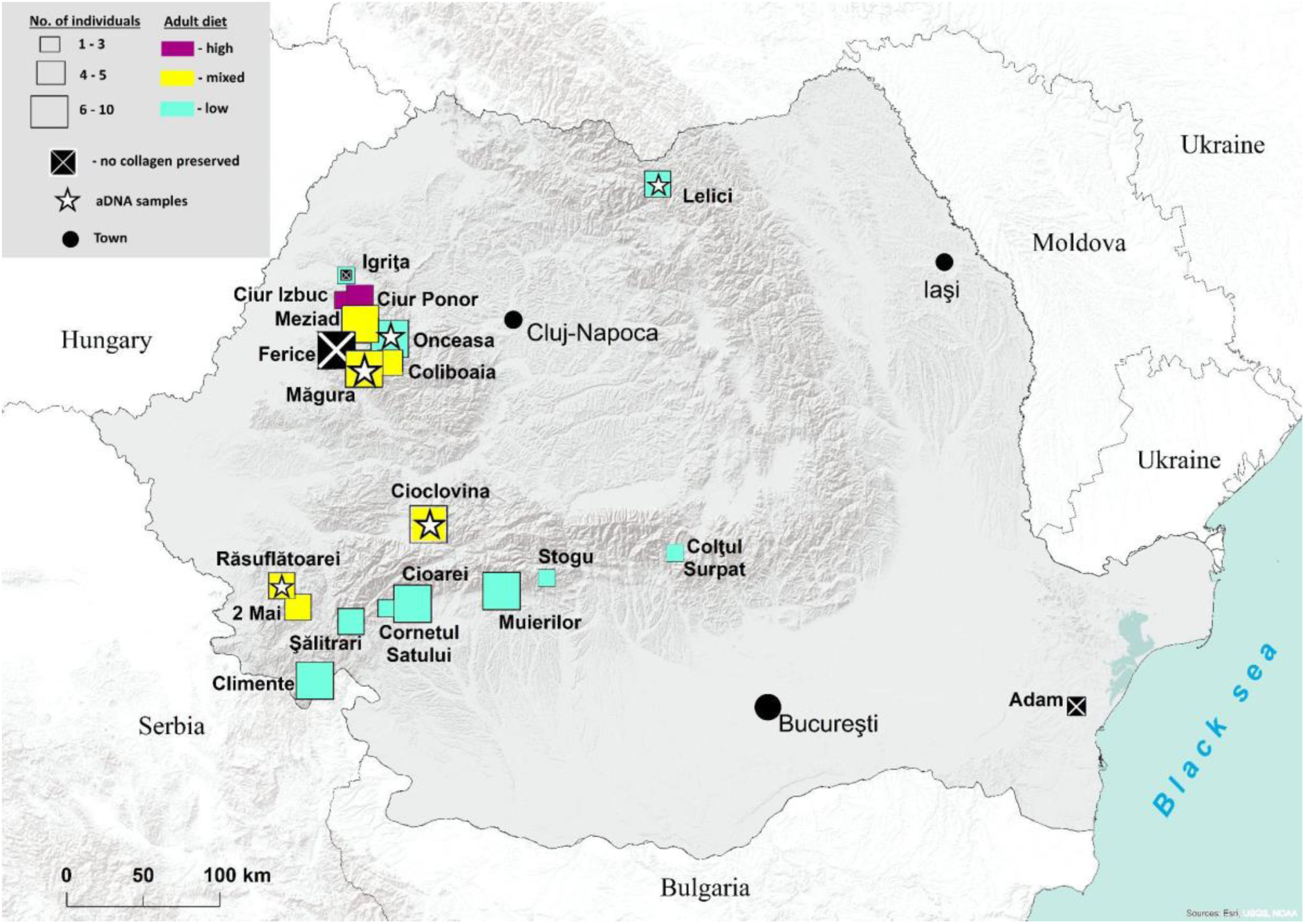
Cave bear sampling sites map. Map with sampling localities showing the number of individuals from each site (square size - small square n < 3; medium-sized square 3 ≤ n < 5; big square 5 ≤ n < 10). Additionally, squares are colour-coded based on the summary of *δ*^15^N isotopic results: low, high, and mixed (both low and high) *δ*^15^N values. Igrița and Șelitrari caves provided both poorly and well-preserved collagen samples. Basemaps sources: World Countries layer: ESRI and Garmin International, Inc., US Central Intelligence Agency, 2021; ESRI Light Grey Canvas map service (http://server.arcgisonline.com/arcgis/services/Canvas/World_Light_Gray_Base/MapServer).

### Palaeodiet analysis based on carbon and nitrogen isotopic composition of bulk collagen

To analyse the diet of Late Pleistocene cave bears of the Romanian Carpathians we studied newly generated and published *δ*^13^C and *δ*^15^N signatures (Naito et al., 2020) from bulk collagen of 105 individuals from 20 caves. The bone collagen extraction was performed following the protocol described by (Bocherens et al., 1997). The carbon and nitrogen stable isotopic measurements were performed using an elemental analyser NC 2500 (Carlo Erba, Milan, Italy) connected to a Thermo Quest Delta + XL mass spectrometer (Thermo Fisher Scientific, Bremen, Germany) (Bocherens, 2014). The reliability of the isotopic fingerprints of the collagen extracts was assessed using their chemical composition by checking whether %C, %N, C/N were well within the values expected for well-preserved collagen by comparison with collagen extracted from fresh bones using the same protocol (Ambrose, 1990; DeNiro, 1985) (Table S1). Isotopic compositions are expressed in ‰ using the *δ* notation. The international reference standards were used for *δ*^13^C values (V-PDB) and *δ*^15^N (atmospheric AIR). Measurements were then normalised to *δ*^13^C values of USGS24 and to *δ*^15^N values of IAEA 305 A. Analytical error, based on one standard-deviation of the mean of multiple analyses of purified collagen from modern bones and international standards was ±0.1 ‰ for *δ*^13^C measurements and ±0.2 ‰ for *δ*^15^N measurements.

Principal component analysis (PCA) was used to explore patterns in isotopic signature (*δ*^13^C and *δ*^15^N) in relation to altitude (quantitative variable) and geographic position (qualitative variable) with respect to the Carpathian Mountains. The analysis was performed and plotted using the packages FactoMineR: Multivariate Exploratory Data Analysis and Data Mining version 2.3 (Lê et al., 2008) and factoextra version 1.0.7 in R (R Core Team, 2018).

### Radiocarbon dating

Cave bear collagen was dated directly using AMS radiocarbon dating. Radiocarbon dating was performed in the Radiocarbon Laboratory of the University of Zurich. The obtained radiocarbon ages of 7 individuals were 2-sigma calibrated using OxCal v4.4.4 (Ramsey, 2017), based on the IntCal20 atmospheric curve (Reimer et al., 2020) (Table S1).

### Palaeogenome sequencing and analysis

#### Ancient DNA extraction and library preparation

Ancient DNA was isolated from 9 petrous bones from 5 different localities, 3 containing both dietary niches, following the protocol of (Dabney et al., 2013) with reduced centrifugation speeds as described in (Basler et al., 2017). Each DNA extract was quantified with the Qubit 2.0 fluorometer with high sensitivity reagents (Thermo Fisher Scientific) using one microlitre of each 25 μL DNA extract, prior to library preparation. Illumina single-stranded libraries were prepared following the protocol of (Gansauge & Meyer, 2013), after the removal of uracil residues and abasic sites by treating the DNA with uracil-DNA glycosylase and Endonuclease VIII enzymes. The optimal number of library amplification PCR cycles was assessed with qPCR as described in (Basler et al., 2017). A volume of 80 μL including 20 μL template library, Accuprime Pfx DNA polymerase and eight base-pair tailed-primers was used for Indexing PCR to generate dual-indexed library molecules. Commercial silica spin-columns (Qiagen MinElute) were used to purify the amplified libraries. Prior to single-end sequencing on Illumina platforms, library concentration and length distribution were quantified using Qubit 2.0 and 2200 TapeStation (Agilent Technologies), respectively.

#### Sequencing and bioinformatic analyses

Shotgun sequencing was performed on an Illumina NextSeq 500 sequencing platform. Based on the procedures described in (Paijmans et al., 2017), 75 bp single-end reads using the NextSeq 500/550 High Output Kit v2 (75 cycles) sequencing kit were obtained. Raw reads were trimmed with minimum overlap of one nucleotide and reads shorter than 30 bp were discarded using cutadapt v1.12 (Martin, 2011). Mapping was performed using bwa v0.7.15 with the “aln” algorithm (Li & Durbin, 2009). For mitochondrial DNA the processed reads were mapped to a reference mitogenome sequence of *Ursus spelaeus* (Genbank Acc. No. EU327344, (Bon et al., 2008)). For nuclear DNA, the processed reads were mapped to an *Ursus arctos* genome assembly (GenBank assembly, GCA_023065955.1). The mapped reads were filtered for mapping quality (Q ≥ 30), sorted by 5’ mapping position, and potential PCR duplicates removed (rmdup) with SAMtools v1.3.1 (Li et al., 2009). The authenticity of the sequences mapped to the brown bear reference genome assembly was then assessed by calculating fragment length distributions and checking for the presence of ancient molecular damage caused by cytosine deamination at the fragment ends, using the program mapDamage 2.0 (Jónsson et al., 2013).

#### Mitochondrial genome reconstruction and alignment

Consensus sequences were generated using ANGSDv0.920 (Korneliussen et al., 2014). Using the option-doFasta 3 that takes into account bases with highest effective depth (Wang et al., 2013), further filtering for read mapping and Phred base quality scores >30. The mitogenomes were then aligned to a published alignment of cave bear mitochondrial sequences (Fortes et al., 2016), including also the two mitogenomes published by (Naito et al., 2020). A repetitive section of the d-loop was removed from the alignment to avoid errors. The final alignment comprised 28 sequences of 16,468 nucleotides. The GenBank accession numbers of the mitogenomes generated in this study are: MW598260 – MW598268.

#### Genome population structure

Nuclear genome analyses were performed in ANGSDv0.920 (Korneliussen et al., 2014) using the following quality filters: a minimum base quality score of 30 (-minQ 30), minimum mapping quality score of 30 (-minMapQ 30), a maximum depth < the 95^th^ percentile of global coverage (-setMaxDepth), transitions excluded (-rmTrans 1), requiring minor alleles to be observed in at least two individuals (-minFreq), and only retaining sites where all individuals had a base (-minInd). Scaffolds below 1 Mb were excluded.

To summarize overall genetic variation among individuals, a Principal Component Analysis (PCA) was performed using single read sampling (-doIBS 1). The resulting covariance matrix was then used for PCA using the princomp() function in R (Figure 2B). An IBS matrix was calculated using the same filters and including an *eremus* cave bear as an outgroup for the phylogenetic reconstruction of cave bears (Figure 2C). The tree was calculated using the neighbor joining clustering method in the R package Analyses of Phylogenetics and Evolution (ape) (Paradis et al., 2004); R Core Team, 2018), rooted using the *eremus* cave bear.

**Figure 2.**
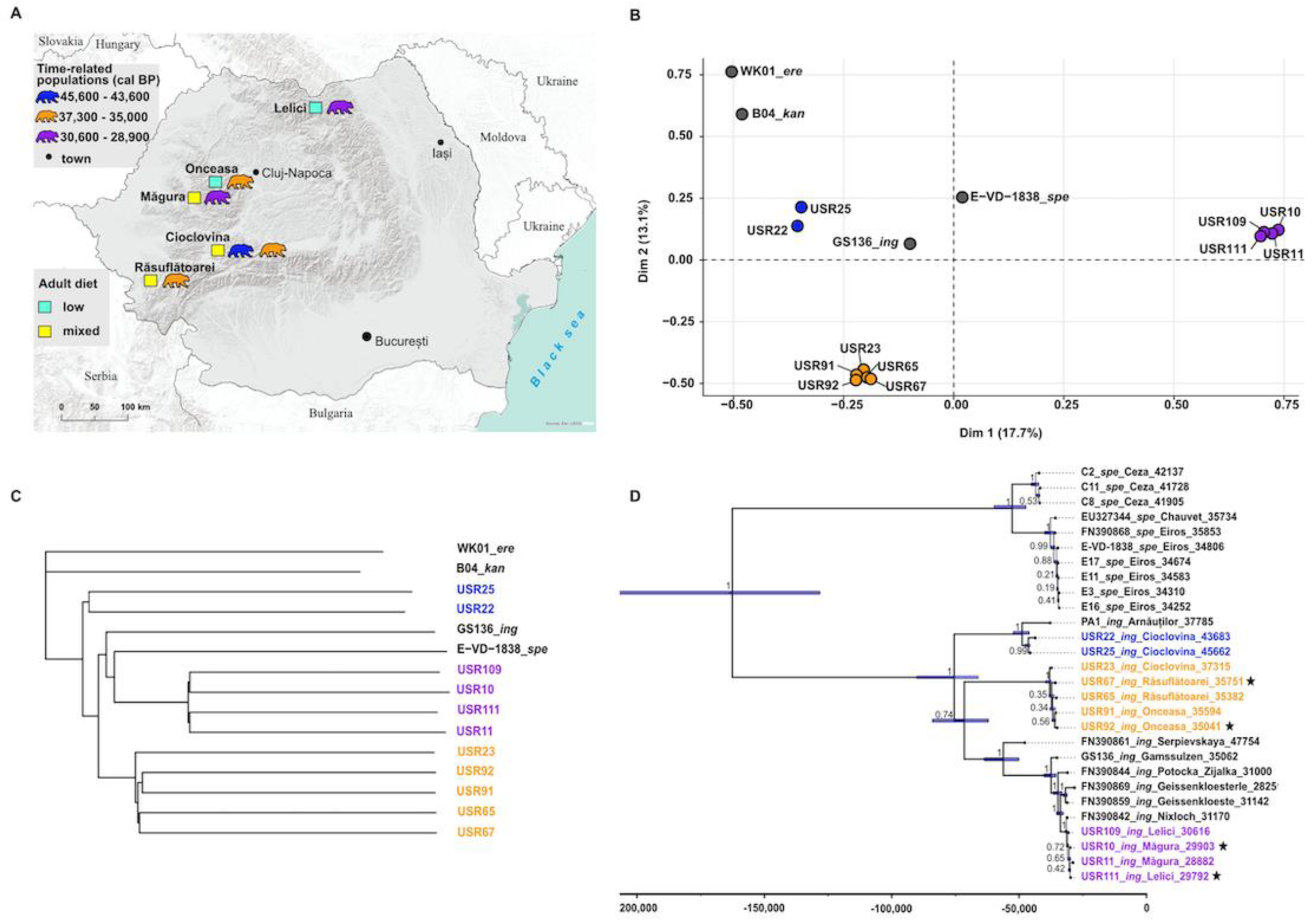
Spatial distribution of time-dependent cave bear populations through palaeogenomic analysis. **A.** Map of sampled localities for aDNA coloured by the inferred time-dependent cave bear populations. Colours indicating inferred adult diets are consistent with those used in Figure 1. **B.** Principal Components Analysis for the Romanian samples (USR) (circles coloured according to the time-structured populations) and other bears from Europe (dark grey circles) from 125,375 transversion sites. Percentages on the X and Y axis represent the percentage of variance explained by each respective component. **C**. Neighbor joining tree based on an IBS matrix of the same nuclear genome dataset. Samples from Romania are coloured according to the identified time-dependent populations. The phylogenetic reconstruction confirmed that each time-dependent population formed a monophyletic group. **D.** Time-calibrated mitochondrial phylogeny of the cave bears. The lower scale shows thousand years BP. Branch labels indicate posterior clade probabilities. Nodes are centred on the median estimated divergence time, and bars show the 95% HPDs. Black stars: estimated age of the sample. USR: code for samples from Romanian Carpathians. Other samples from (Fortes et al., 2016). USR10 and USR67 are from (Naito et al., 2020).

#### Genetic sex determination

The sex of each sample was determined by analysing reads mapped to both the X-chromosomes and autosomes (36) and calculating the X-chromosome to autosome ratio. In the context of completely random shotgun sequencing, an expected normalized ratio of approximately 0.5 suggests males, while ratios of approximately 1.0 indicate females (Figure S3). Pearson’s Chi-squared test with Yates’ continuity correction was conducted to evaluate potential differences between females and males in dietary patterns.

#### Molecular tip-dating

Ages of four samples were estimated using Bayesian phylogenetic tip dating, which has proven to be reliable for Late Pleistocene European cave bears (Fortes et al., 2016) (Table S2). Data partitioning and substitution model selection was performed using PartitionFinder v.2.1.1 (Lanfear et al., 2012) (Table S3). All substitution models available in BEAST v.1.8.2 were considered, and the best partitioning scheme was computed under the Bayesian Information Criterion using linked branch lengths and the greedy search algorithm (Drummond et al., 2012) Separate tip-dating analyses were performed for 4 individuals without radiocarbon dates, using the median calibrated ^14^C dates of dated individuals as calibration (from the dataset in (Fortes et al., 2016; Naito et al., 2020). A uniform, uninformative prior of zero to one million years BP was used to sample the posterior distribution for each tip date. A piecewise-constant coalescent Bayesian Skyline population model was selected to accommodate changes in effective population size during the time-span of the tree. Unlinked strict molecular clocks were set for each data partition, with the posterior distribution of the substitution rate of each sampled with a uniform, uninformative prior of zero to 20% per million years. To assess if the MCMC chain had run for a sufficient length to achieve burn-in and adequate sampling of all parameters (ESS > 200) the software Tracer v1.6 was used (Rambaut et al., 2018). TreeAnnotator was used to select the Maximum Clade Credibility Tree from the posterior sample, which was visualised in FigTree v.1.4.4 (http://tree.bio.ed.ac.uk/software/figtree/) (Figure 2D).

#### Vegetation modelling maps

We utilised the palaeovegetation dataset provided by (Allen et al., 2020) and employed the inverse distance weighted tool (IDW, ESRI ArcMap 10.8.) to interpolate the carbon concentration of boreal trees and boreal shrubs within European countries hosting cave bear localities dated from MIS 3 to MIS 2 (Figure 4, Dataset S2). The interpolation was calculated for the statistical mean of the following time intervals: 46 - 44 ka BP, 38 - 36 ka BP and 30 - 28 ka BP, intervals corresponding to the ages of studied cave bear populations. Boreal tree carbon concentrations were mapped from low (under 3.5 billion kg carbon) to very high values (over 15.5 billion kg carbon). Similarly, we mapped the boreal shrub concentrations using intervals from low (under 500 million kg carbon) to very high values (7 billion kg carbon). We also mapped the extent of the European ice sheet for each interval using data from the same dataset (Allen et al., 2020).

## Results

### Spatial patterns of dietary variation

Bulk bone collagen of 95 adult individuals from 18 caves showed good collagen preservation (Figure 1, Table S1). As previously observed, the *δ*^13^C and *δ*^15^N signatures of cave bears from the Romanian Carpathians show a greater variability compared to cave bears from the rest of Europe (Austria, Belgium, France, Germany, Italy, Poland, Slovakia, Slovenia, Spain, Switzerland) (Figure S2). The highest variability in *δ*^13^C and *δ*^15^N values was found between individuals sampled from caves located within the intra-Carpathian basin, with both isotopic extremes frequently occurring within the same cave (Figures S2 and S3). Principal Component Analysis (PCA) was conducted on *δ*^13^C and *δ*^15^N values, as well as the altitude of the caves of provenance as quantitative variables, while considering the geographical pattern as a qualitative variable. Examination of the first 2 PCA axes (which explain 81% of the total variation in the dataset) showed that low (< 100 m a.s.l.) and high (> 1,000 m a.s.l.) altitudes are associated with the clustering of individuals with low *δ*^15^N values in the lower left part of the plot (all samples from Climente I), and in the upper right part of the plot, respectively (all samples from Onceasa and Cave 7 in Lelici referred to as Lelici hereafter). Individuals with high *δ*^15^N values located within the intra-Carpathian basin, at altitudes < 1,000 m a.s.l., clustered in the lower right part of the plot (Figure S3).

### Population structure

We investigated the structuring of cave bear populations and its relation to dietary variation by analyzing DNA extracted from fossil petrous bones of eleven individuals collected from five of the sampled caves (Figure 2A).

PCA results, based on 125,375 informative genomic sites, revealed the presence of three distinct populations (Figure 2B). Based on radiocarbon dates of seven of the eleven investigated individuals, the oldest population dates to approximately 45,000 years before present (45 ka BP) and includes two individuals from Cioclovina cave. The middle-aged population, at approximately 36 ka BP, encompasses five individuals from three caves: Cioclovina, Onceasa, and Răsuflătoarei. The youngest population, aged around 29 ka BP, consists of four individuals from two caves: Măgura and Lelici. The inferred population structure shows no obvious relationship with their spatial distribution: individuals from different caves and even massifs fall within the same cluster when they occur in the same time period (Figure 2B and 2C). We similarly find no obvious relationship between population structure and diet, as both dietary niches co-occur within all three time-dependent populations.

### Molecular dating

We further assessed temporal population structure by dating the four individuals that lacked radiocarbon dates using molecular phylogenetic tip dating based on mitochondrial genome sequences, with a set of radiocarbon dated sequences providing time calibration. This analysis further supported the occurrence of three distinct genetic clusters among individuals from the Romanian Carpathians, dating to approximately 45 ka, 36 ka, and 29 ka BP, respectively (Figure 2D). These clusters form separate mitochondrial clades that correspond with the inferred whole-genome population structure. These three clades are not reciprocally monophyletic, as the young cluster is nested closely within a clade comprising other *ingressus*

European cave bears, reinforcing their genetic distinctiveness.

In conclusion, neither nuclear nor mitochondrial markers revealed any correlation between spatial origin of the samples and/or temporal population structuring and the two dietary niches.

### Sex identification

Sex assignment was carried out both morphologically and/or genetically for individuals with genome sequencing, based on the ratio of the number of reads mapped to X chromosome versus autosomes (Figure S3). These analyses revealed no predisposition for isotopic variability between the sexes, both within individual caves and across multiple caves, as indicated in Table 1. Notably, variability in isotopic values was observed in both males and females. Out of 16 males, 4 displayed high *δ*^15^N values (comprising 25%), while out of 21 females, 9 displayed high *δ*^15^N values (accounting for 43%). The Chi-squared test with Yates’ continuity correction indicates no statistically significant difference in the frequencies of isotopic pattern observations between the female and male groups (X-squared = 0, p-value = 1).

## Discussion

Our results reject time period, geographic location, sex, and population affinity as the principal factors driving dietary variation in Romanian cave bears. We find that bears from the same population, living within the same time span (i.e., ∼ 45-43 ka BP; ∼ 37-35 ka BP; ∼ 30-29 ka BP) and utilising the same hibernation spaces, exhibited distinct dietary preferences. Moreover, these dietary differences persisted over the adult lifetimes of the animals. This individual variation suggests the presence of learning behaviour, as it implies that bears adjusted their feeding habits based on personal experiences rather than genetic or environmental factors alone.

(Kyriazakis et al., 1999) highlight that dietary choices in natural environments are largely influenced by the constantly changing availability and composition of food sources, affected by both environmental and social factors. This variability leads animals to select diets tailored to their specific needs, underscoring the importance of learning in the plasticity of feeding behaviour. Learning equips animals to respond to both temporal and spatial shifts in their feeding environments. Complementing this view, (Roper, 1986) proposed a framework for understanding this plasticity. According to Roper, the evolution of plasticity in feeding behaviour relies on two primary factors: the underlying drivers of behavioural variations, and the methods through which these alternative behaviours are passed from one individual to another. This approach weaves together the concepts of environmental influence and individual learning, providing a comprehensive view of how animals adjust their feeding strategies over time.

### Drivers of foraging behavioural plasticity

The dietary differences observed in cave bears inhabiting the mid-altitude regions of the Romanian Carpathians (i.e., between 450 - 770 m a.s.l) may be linked to specific ecosystems that provided the opportunity for dietary niche specialisation and divergence. The more negative *δ*^13^C values exhibited by some Romanian cave bears are typically associated with densely forested environments, which coupled with slight difference in *δ*^13^C values between individuals, suggest that these populations inhabited forest ecosystems characterized by similar environmental conditions and available plant resources (Ambrose & DeNiro, 1989). Considering dietary flexibility observed within the same trophic level, particularly herbivory (Ambrose & DeNiro, 1989; Naito et al., 2020), at mid-altitudes, it may be linked to the consumption of plants with varying *δ*^15^N signatures within forest ecosystems. This is supported by significant variations in *δ*^15^N values among herbivores in the same ecological context, attributed to their consumption of diverse plant categories across Late Pleistocene regions, as shown in previous studies (e.g., (Drucker et al., 2015; Jürgensen et al., 2017). Bocherens (2019) provided a comprehensive overview of nitrogen sequestration variations in different plant species, resulting in substantial disparities in their *δ*^15^N signatures. Shrubs and trees are associated with lower *δ*^15^N values (-5‰ to +5‰), while various grasses, sedges, forbs, ferns and mushrooms exhibit elevated *δ*^15^N values (-2‰ to +9‰) (Gundale et al., 2012; Hobbie et al., 2005).

The palaeoclimate and palaeovegation records from Europe show that central and western Europe were covered by an extended tundra-steppe biome with less developed forest cover during the Late Pleistocene (Moreno et al., 2014). In contrast, the Carpathian region in eastern central Europe has been subjected to lower amplitude cooling and drying during this time period (Feurdean et al., 2014; Staubwasser et al., 2018). The less pronounced climate fluctuations in the Romanian Carpathians led to the presence of extensive boreal forest ecosystems, comprising species such as *Pinus, Betula, Acer, Alnus, Salix*, along with small patches of temperate deciduous trees including *Ulmus, Quercus,* and *Carpinus*. These ecosystems were particularly developed during warmer periods (interstadials) of the Late Pleistocene. Therefore, it is plausible that diverse forest compositions in the Romanian Carpathians have facilitated a more varied plant diet than in central and western Europe. Variation in forest cover likely contributed to higher spatial variability in *δ*^15^N signatures for cave bear forage plants, with areas characterised by sparse or denser forest cover, exhibiting low and high *δ*^15^N values, respectively.

The cave bear populations dated to 45-43 ka BP and 37-35 ka BP are associated with warmer conditions and the presence of extensive boreal woodlands, as inferred from proxy-based reconstructions and simulations (Figures 3, 4). Conversely, the youngest population, estimated to be around 29 ka BP, corresponds to a cold and dry climatic event of the Late Pleistocene [Marine Isotope Stage (MIS)2, Heinrich (H) 3 stadial], characterised by extended steppe-tundra vegetation also in the Romanian Carpathians (Feurdean et al., 2014, 2015; Pendea et al., 2009; Staubwasser et al., 2018). Limited proxy-based records of Late Pleistocene vegetation from Romania, restricts our ability to determine spatial and temporal variability in plant resources and dietary niche variation in cave bears. However, vegetation modelling generally supports the reconstructions by suggesting that the two older time periods (45-43 ka BP, 37-35 ka BP) had a higher boreal tree biomass than the youngest period (30-29 ka BP) (Figure 4). Additionally, modelling results suggest that tree biomass was greater in the southwest Romanian Carpathians during the oldest periods (46-44 ka BP), and that there was less spatial variability in tree biomass during 38-36 ka BP, and higher tree biomass in the Apuseni Mountains than in the southwest Romanian Carpathians from 30-28 ka BP.

**Figure 3.**
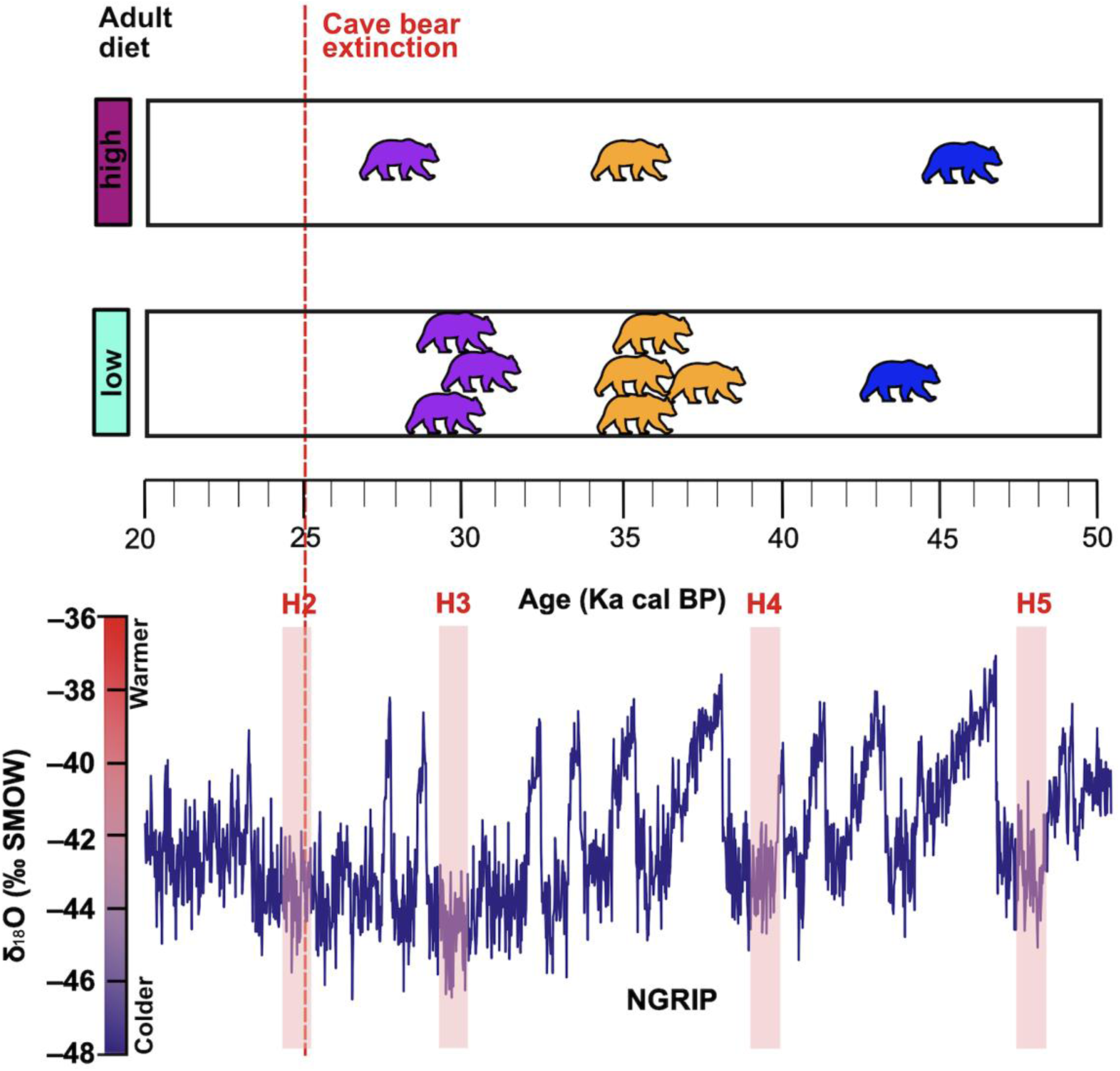
Timeline indicating the three populations related to the adult diet and past climatic changes. Cave bears, representing individuals from three time-dependent populations, are coloured based on their inferred time-dependent populations of Romanian Carpathians. Colours indicating inferred adult diets are consistent with those used in Figure 1. NGRIP, dark blue curve: North Greenland Ice Core Project (NGRIP) *δ*^18^O (Rasmussen et al., 2014), including Heinrich events (H) (Hemming, 2004). Bears are centered on the inferred median age.

**Figure 4.**
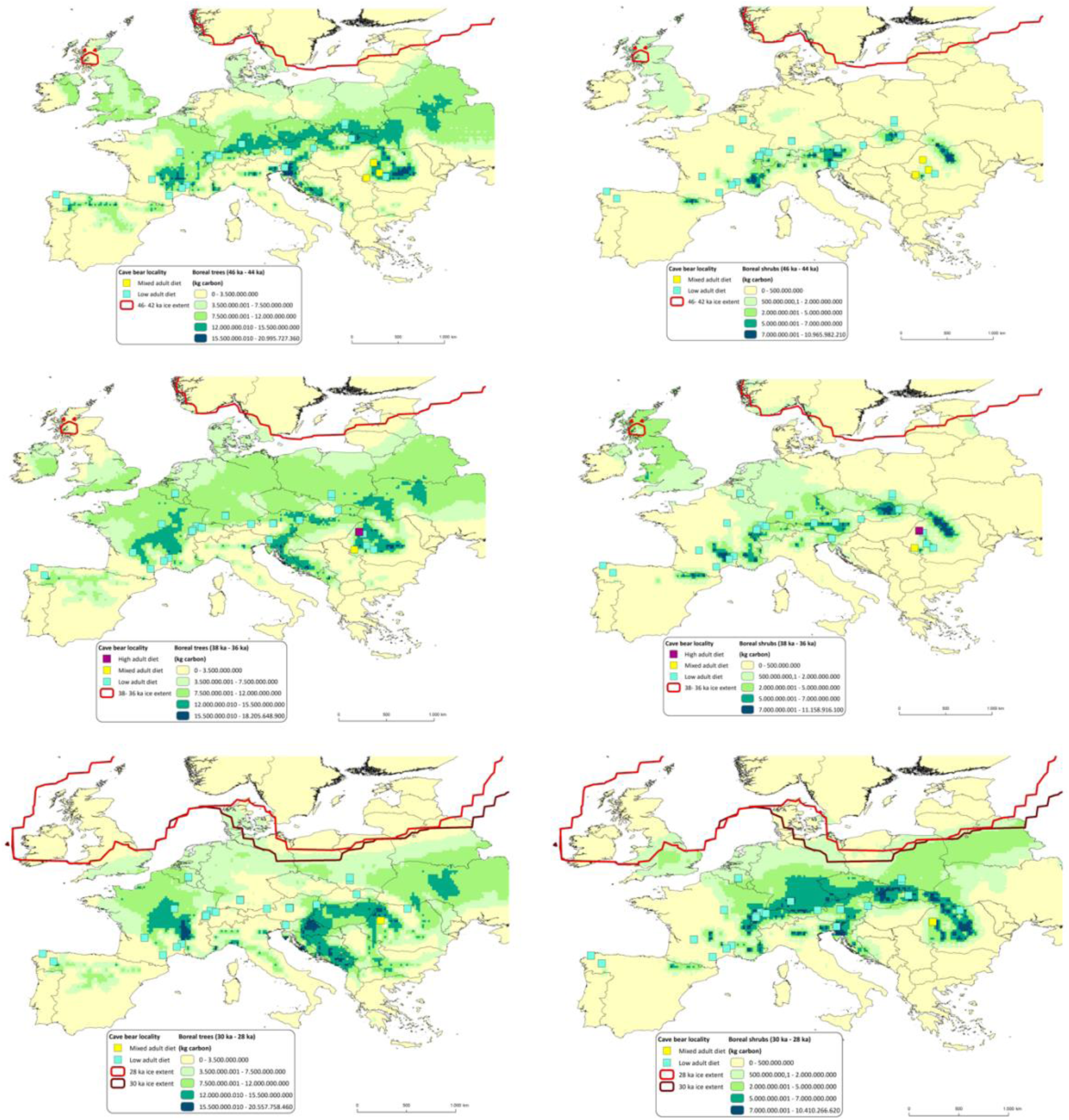
Palaeovegetation modelling scenarios for the distribution of boreal trees (left panels) and shrubs (right panels) based on the three inferred time periods of Romanian cave bear populations (rows). The cave bear localities from Europe in each scenario correspond to cave bear occupation dating from MIS 3 to MIS 2. Colours representing the localities indicate inferred adult diets and are consistent with those used in Figure 1. The colour ramp from yellow to dark green for palaeovegetation layer matches the low to very high concentrations of boreal trees and shrubs expressed in kg carbon.

These patterns in boreal tree biomass contrast with those observed in other Romanian regions, as well as in Central and Western Europe. Furthermore, the consistency of these vegetation patterns with the observed variability in isotopic signatures suggests that cave bears sampled in these areas had access to more diverse plant resources, which may have promoted the development of unique dietary niches within the same geographic area.

An additional factor that may have influenced dietary niche divergence among cave bears is competition, potentially both intraspecific, and interspecific competition with the sympatric brown bear. Given the very large cave bear populations in the area, as documented by (Robu, 2015), intraspecific dietary partitioning within the same trophic niche might have occurred, as documented in species like water snakes (Perkins et al., 2020) and bats (Mata et al., 2016). This partitioning would minimise competition by diversifying food preferences and expanding niches. Furthermore, interspecific competition with sympatric brown bears may have additionally driven dietary shifts towards alternate plant sources. This concept aligns with findings by (Belant et al., 2006), who demonstrated that in sympatry, ursid species, especially brown bears, can influence American black bears to alter their diet, often resulting in the consumption of less nutritious food sources. This indicates a pattern of dietary flexibility among bear species in response to competitive pressures in shared habitats.

### Mechanisms of foraging behavioural plasticity

Unlike findings from other cave bear populations, where the segregation of individuals into exclusive mitochondrial clades was associated with dietary differences, supported by minor yet significant isotopic variations in regions like the Austrian Alps (Bocherens et al., 2011) or correlated with specific hibernation caves in cave bears from northern Spain (Fortes et al., 2016), our results diverged from these patterns. Phylogenetic lineage diversification was unrelated to niche variation or geographical location. Intrapopulation niche variation in bears might be explained by social learning (cultural inheritance) and/or asocial learning for niche partitioning, as evidenced in different living bear species. For instance, in American black bears, social learning has been documented as the primary mechanism responsible for foraging on human foods in Sequoia and Yosemite National Parks (Hopkins, 2013; Mazur & Seher, 2008). Similarly, in brown bears, learning has been linked to conflict behaviour and habitat selection, as detailed by (Morehouse et al., 2016). Additionally, their diet selection is closely tied to individual experiences after they become independent from their mothers. In heterogeneous and challenging environments like the Arctic, brown bears exhibit significant dietary variation influenced by time-dependent spatial memory, indicating complex cognitive processes in locating resource-rich food patches (Thompson et al., 2022). In the case of polar bears, learning has been associated with adapting to changing sea ice conditions on an annual basis, showing behavioural plasticity in on-shore/off-shore activities (Lillie et al., 2018).

This individual dietary specialisation is influenced by several factors, including habitat exploration, the acquisition of foraging experience, and, in the case of males, social dominance. Interestingly, female brown bears tend to maintain the average diet of their natal habitat, as highlighted in the research by (Jimbo et al., 2022). Our research revealed that there was no clear difference in the *δ*^15^N variability of the diets between male and female cave bears. Both sexes exhibited variations in *δ*^15^N levels, both within individual caves and across different caves. Given that the Pearson’s Chi-squared test with Yates’ continuity correction showed no significant differences between males and females, it’s important to interpret the observed trends cautiously, considering potential variations within the dataset. While our data did not show clear distinctions between the sexes, it is worth noting that we observed a greater number of female bears with higher *δ*^15^N values. This might suggest that female cave bears tended to remain more consistent with their original habitats, in contrast to male cave bears who tended to wander more extensively. Broader roaming behaviour of male cave bears would have given them access to a more varied selection of plants, leading to a higher proportion of males exhibiting lower *δ*^15^N values in their dietary profile.

American black bears exhibit well-documented instances of learning facilitated by close kin relationships, such as mother-to-offspring transmission, and cultural inheritance (Hopkins, 2013). In killer whales, foraging specialisation appears to be passed down generationally through mimicry and learning (Ford et al., 2010). Moreover, these foraging strategies significantly influence their mitochondrial genetic structure, resulting in foraging specialists within ecotypes sharing similar diets, while differences in diet and genetic composition increase with geographical distance within each ecotype (Hoelzel et al., 2007). Despite the lack of an apparent correlation between mitochondrial genetic structure and diet in our study, the possibility of mother-to-offspring transmission cannot be ruled out. This mechanism may have occurred, although not visibly reflected in the observed genetic patterns, possibly due to the varying number of generations required for completion, contingent upon population sizes. However, it is possible that individual foraging behaviours are influenced more by learning experiences, leading to the development of individualistic foraging strategies, possibly based on early foraging success after cubs become independent from their mothers. This behaviour is then maintained throughout their lives. An alternative explanation could be a creche-based social system for offspring care, similar to what (Sheppard et al., 2018) observed in mongooses, where such a system resulted in a decoupling of isotope signature and genetic relatedness. Additionally, as (Murie, 2012) demonstrated in grizzly bears, behavioural flexibility can occur in dense, short-term aggregations in areas rich in food, where females with young may associate and cooperatively care for and nurse the young. This suggests that there might have been a complex and varied learning process in cave bears, involving both individual experiences and potentially communal learning mechanisms.

## Conclusion

Our findings indicate that the distinct dietary niches observed in Romanian cave bears do not separate according to specific time periods, geographic location, sex or population affinity. Instead, both dietary niches co-occur across each of these potential explanatory variables. Dietary variation is therefore most likely attributed to individual behavioural responses to the availability of food resources, potentially relating to changes in forest cover. A changing environment may force individuals to learn new strategies and possibly make new associations. In the context where isotope values reflect an organism’s dietary patterns throughout its adult lifetime, our findings suggest that some individuals exhibited a tendency to explore new food sources beyond the standard dietary options observed in most European cave bears. This led to variations in their food intake, even though they occupied the same overall herbivorous niche. These results provide compelling evidence of the flexible capacity and resilience of these individuals over their lifespan. Guided by positive experiences related to specific dietary choices, they learned to consume differently compared to most of their conspecifics.

The interdisciplinary method used in this study unlocks a deeper understanding of behavioural processes, revealing details that would remain hidden if each type of evidence were examined separately. With the recent advancements in exploring DNA-environment interactions through ancient epigenomes, future research should concentrate on investigating the influence of epigenetic mechanisms in behavioural plasticity. This direction could provide a more comprehensive understanding of how extinct species adapted to their environments.

## Acknowledgements

The authors thank Viorel Traian Lascu; Bogdan Bădescu, Iosif Morac and Exploratorii Reșița Caving Club; Adrian Done and Bucovina Caving Club; Tudor Rus and Speodava Caving Club, and Prusik Caving Club for help during fieldwork. We are grateful to Daniel Veres for providing insights in earlier versions of the manuscript. We would also like to thank the administrations of the Apuseni Natural Park, and Semenic-Cheile Carașului and Domogled Valea Cernei National Parks for access to some of the investigated caves and the National Museum of Natural History “Grigore Antipa’’ for access to their fossil collection. We acknowledge the support of the Supercomputing Wales project, which is part-funded by the European Regional Development Fund (ERDF) via the Welsh Government. This work was financially supported by the Alexander von Humboldt Foundation, grant number ROU 1161809 HFST-P to INM. During the preparation of this paper INM was financed through the European Union’s Horizon 2020 research and innovation programme under the Marie Skłodowska-Curie grant agreement No 885088.

## Statement of authorship

Conceptualisation, I.N.M., M.H., A.B., and H.B.; Methodology, I.N.M., M.N.-A., A.F., M.H., A.B., and H.B.; Molecular work, I.N.M. and F.A.; Isotope work: I.N.M. and D.G.D.; Sampling: I.N.M., M.R., and M.V.; Writing - Original draft, I.N.M with input from A.B., M.H., H.B., M.N-A., A.F.; Writing – Review & Editing, all authors; Resources, I.N.M., A.B., M.C., E.C.N., M.H., A.B., and H.B; Supervision, M.H., H.B., and A.B; Project Administration: I.N.M., M.H., and H.B.; Funding Acquisition, I.N.M. All authors gave final approval for publication.

## Data accessibility

DNA mitochondrial sequences obtained in this study are deposited in GenBank with accession numbers: MW598260 – MW598268. Raw sequence reads will be deposited in the EMBL Nucleotide Sequence Database (ENA) upon acceptance under the accession numbers: XXX. All isotope data and palaeogenomic mapping data supporting the tables and figures are provided in the Supporting Information.

