## supplements for "Ancient biomolecules suggest learned foraging strategy in extinct cave bears"

### Supporting information

Supporting information includes two datasets, three tables and three figures.

**Dataset S1.** Palaeogenomic dataset including number of reads, reads mapped to brown bear, depth of coverage and endogenous content.

**Dataset S2.** Palaeodiet dataset for European and Romanian cave bears, incorporating data from published sources as well as samples from this study dated from MIS 3 to MIS 2. This dataset was used to generate Figure 4.

**Table S1.** $\delta^{13}C$ and $\delta^{15}N$ signatures from bulk collagen of the Romanian Carpathian cave bears. In red: samples not included in downstream analyses due to poor collagen preservation or badly preservation making collagen extraction impossible. * from Naito et al. (2020). # genetic sex assignment

| $\delta^{13}C$ and $\delta^{15}N$ signatures from bulk collagen | | | | | | | |
|---|---|---|---|---|---|---|---|
| **Sample ID** | **Sampling cave** | **% C collagen** | **% N collagen** | **C / N atom** | **$\delta^{13}C$ (‰) (VPDB)** | **$\delta^{15}N$ (‰) (AIR)** | **Sex#** |
| USR1 | Meziad | 36.9 | 12.9 | 3.3 | -20.5 | 1.7 | |
| USR2 | Meziad | 39.3 | 14.0 | 3.3 | -21.1 | 6.2 | |
| USR3 | Meziad | 23.2 | 8.2 | 3.3 | -20.9 | 5.6 | |
| USR4 | Meziad | 37.8 | 13.3 | 3.3 | -21.1 | 6.3 | |
| USR5 | Meziad | 39.4 | 14.0 | 3.3 | -20.7 | 5.5 | |
| USR6 | Meziad | 28.3 | 10.0 | 3.3 | -20.8 | 5.4 | |
| USR7 | Meziad | 35.9 | 12.7 | 3.3 | -21.5 | 5.8 | |
| USR8 | Meziad | 34.0 | 12.0 | 3.3 | -21.3 | 4.1 | |
| USR9 | Meziad | 35.7 | 12.5 | 3.3 | -21.9 | 8.1 | |
| USR10* | Măgura | 26.0 | 9.1 | 3.3 | -22.0 | 6.3 | F# |
| USR11* | Măgura | 36.8 | 12.3 | 3.5 | -22.1 | 8.1 | F# |
| USR12 | Măgura | 38.8 | 13.6 | 3.3 | -21.1 | 6.2 | |
| USR13 | Măgura | 42.5 | 14.9 | 3.3 | -20.9 | 5.3 | |
| USR14 | Măgura | 35.8 | 12.4 | 3.4 | -21.2 | 4.7 | |
| USR15 | Măgura | 37.2 | 13.1 | 3.3 | -21.2 | 5.7 | |
| USR16 | Măgura | 36.4 | 12.7 | 3.4 | -21.3 | 6.0 | |
| USR17 | Măgura | 35.2 | 12.0 | 3.4 | -22.3 | 7.1 | |
| USR18 | Măgura | 35.4 | 12.3 | 3.4 | -21.1 | 6.1 | |
| USR19 | Măgura | 32.1 | 11.1 | 3.4 | -21.4 | 5.3 | |
| USR20 | Cioclovina | 40.4 | 14.2 | 3.3 | -21.7 | 6.7 | |
| USR21 | Cioclovina | 39.3 | 13.6 | 3.4 | -22.9 | 7.3 | |
| USR22* | Cioclovina | 33.9 | 11.8 | 3.3 | -22.0 | 5.2 | M# |
| USR23 | Cioclovina | 34.7 | 12.2 | 3.3 | -22.5 | 3.6 | M# |
| USR24 | Cioclovina | 36.1 | 12.6 | 3.3 | -22.9 | 4.8 | |
| USR25* | Cioclovina | 36.7 | 12.9 | 3.3 | -22.3 | 8.9 | F# |
| USR26 | Cioclovina | 42.7 | 15.0 | 3.3 | -22.6 | 4.8 | F |
| USR27 | Igrița | 30.5 | 10.7 | 3.3 | -21.6 | 3.0 | |
| USR28 | Igrița | 15.6 | 5.5 | 3.3 | -21.0 | 3.9 | |
| USR29 | Igrița | 19.5 | 6.8 | 3.3 | -21.8 | 3.3 | |
| USR30 | Climente I | 43.2 | 15.4 | 3.3 | -20.8 | 2.7 | |
| USR31 | Climente I | 41.4 | 14.6 | 3.3 | -20.8 | 4.5 | |
| USR32 | Climente I | 41.2 | 14.6 | 3.3 | -20.9 | 4.8 | |

| | | | | | | | |
|---|---|---|---|---|---|---|---|
| USR33 | Climente I | 38.1 | 13.2 | 3.4 | -20.6 | 2.5 | |
| USR34 | Climente I | 33.5 | 11.9 | 3.3 | -21.4 | 2.0 | |
| USR35 | Climente I | 41.8 | 14.8 | 3.3 | -20.7 | 2.1 | |
| USR36 | Climente I | 42.3 | 14.9 | 3.3 | -21.1 | 4.1 | |
| USR37 | Climente I | 39.3 | 13.6 | 3.4 | -20.9 | 3.3 | |
| USR38 | Climente I | 41.1 | 14.2 | 3.4 | -21.6 | 1.3 | |
| USR39 | Climente I | 41.3 | 14.3 | 3.4 | -21.3 | 3.8 | |
| USR40 | Şălitrari | 31.8 | 10.2 | 3.6 | -20.6 | 1.9 | |
| USR41 | Şălitrari | 42.6 | 15.1 | 3.3 | -20.8 | 2.1 | |
| USR42 | Şălitrari | 39.5 | 12.1 | 3.8 | -21.7 | 3.9 | |
| USR43 | Şălitrari | 32.9 | 9.7 | 4.0 | -21.2 | 2.3 | |
| USR44 | Colţul Surpat | 39.7 | 13.4 | 3.5 | -23.3 | 5.1 | |
| USR45 | Colţul Surpat | 42.3 | 14.2 | 3.5 | -22.3 | 5.4 | |
| USR46 | Colţul Surpat | 41.6 | 13.4 | 3.6 | -21.3 | 4.2 | |
| USR47 | Stogu | 40.7 | 13.8 | 3.4 | -21.7 | 3.4 | |
| USR48 | 2 Mai | 37.6 | 13.1 | 3.4 | -21.6 | 7.3 | M |
| USR49 | 2 Mai | 41.5 | 14.6 | 3.3 | -21.4 | 4.8 | |
| USR50 | 2 Mai | 41.5 | 14.5 | 3.3 | -21.3 | 4.5 | F |
| USR51 | 2 Mai | 33.2 | 11.2 | 3.5 | -22.3 | 10.9 | |
| USR52 | 2 Mai | 36.9 | 12.7 | 3.4 | -21.3 | 3.1 | |
| USR53 | Adam | 40.5 | 11.0 | 4.3 | -22.8 | 8.2 | |
| USR54 | Adam | 33.0 | 10.3 | 3.7 | -20.9 | 8.0 | |
| USR55 | Muierilor | 32.7 | 11.2 | 3.4 | -21.0 | 2.9 | |
| USR56 | Muierilor | 36.5 | 12.8 | 3.3 | -20.9 | 2.7 | |
| USR57 | Muierilor | 34.3 | 11.9 | 3.4 | -21.4 | 6.7 | |
| USR58 | Muierilor | 40.7 | 14.1 | 3.4 | -20.7 | 3.7 | |
| USR59 | Muierilor | 33.6 | 11.2 | 3.5 | -20.8 | 1.1 | |
| USR60 | Muierilor | 39.8 | 14.1 | 3.3 | -20.8 | 2.1 | |
| USR61 | Muierilor | 33.7 | 11.8 | 3.3 | -20.8 | 2.5 | |
| USR62 | Muierilor | 42.7 | 15.0 | 3.3 | -21.1 | 4.9 | |
| USR63 | Muierilor | 40.4 | 14.4 | 3.3 | -20.7 | 5.3 | |
| USR64 | Răsuflatoarei | 20.0 | 6.9 | 3.4 | -18.5 | 6.3 | M |
| USR65* | Răsuflatoarei | 38.3 | 13.5 | 3.3 | -21.8 | 9.8 | F# |
| USR66 | Răsuflatoarei | 42.7 | 15.1 | 3.3 | -21.6 | 6.5 | F |
| USR67* | Răsuflatoarei | 46.1 | 16.2 | 3.3 | -19.9 | 5.4 | F# |
| USR68 | Răsuflatoarei | 40.1 | 14.3 | 3.3 | -22.4 | 3.5 | M |
| USR69 | 10 din Cornetu Satului | 42.2 | 14.8 | 3.3 | -20.7 | 3.1 | |
| USR70 | 10 din Cornetu Satului | 42.7 | 15.0 | 3.3 | -20.4 | 1.7 | |

| | | | | | | | |
|---|---|---|---|---|---|---|---|
| USR71 | 10 din Cornetu Satului | 37.8 | 13.2 | 3.3 | -21.1 | 2.1 | |
| USR72 | Cioarei de la Boroşteni | 35.8 | 12.4 | 3.4 | -21.2 | 0.8 | |
| USR73 | Cioarei de la Boroşteni | 41.3 | 14.5 | 3.3 | -21.2 | 1.0 | |
| USR74 | Cioarei de la Boroşteni | 37.7 | 13.2 | 3.3 | -21.5 | 2.1 | |
| USR75 | Cioarei de la Boroşteni | 36.4 | 12.9 | 3.3 | -20.4 | 2.8 | |
| USR76 | Cioarei de la Boroşteni | 36.4 | 13.0 | 3.3 | -18.4 | 3.2 | |
| USR77 | Cioarei de la Boroşteni | 39.9 | 14.1 | 3.3 | -18.4 | 6.7 | |
| USR78 | Ciur Ponor | 38.8 | 13.7 | 3.3 | -22.5 | 8.0 | F |
| USR79 | Ciur Ponor | 37.5 | 13.4 | 3.3 | -22.1 | 8.7 | M |
| USR80 | Ciur Ponor | 40.7 | 14.5 | 3.3 | -22.3 | 10.0 | F |
| USR81 | Ciur Ponor | 36.8 | 13.1 | 3.3 | -22.4 | 9.1 | M |
| USR82 | Ciur Izbuc | 26.1 | 9.2 | 3.3 | -18.0 | 7.9 | F |
| USR83 | Ciur Izbuc | 36.1 | 12.7 | 3.3 | -22.7 | 10.6 | F |
| USR84 | Coliboaia | 24.1 | 8.6 | 3.3 | -21.8 | 9.8 | F |
| USR85 | Coliboaia | 36.6 | 13.0 | 3.3 | -21.9 | 8.6 | M |
| USR86 | Coliboaia | 37.2 | 13.2 | 3.3 | -21.9 | 4.2 | F |
| USR87 | Coliboaia | 33.8 | 12.0 | 3.3 | -21.9 | 7.1 | F |
| USR88 | Ferice | - | - | - | - | - | |
| USR89 | Ferice | - | - | - | - | - | |
| USR90 | Ferice | - | - | - | - | - | |
| USR91 | Onceasa | 40.9 | 14.4 | 3.3 | -21.3 | 4.1 | F# |
| USR92 | Onceasa | 41.7 | 14.7 | 3.3 | -22.0 | 5.8 | M# |
| USR93 | Onceasa | 42.6 | 15.2 | 3.3 | -21.3 | 2.5 | F |
| USR94 | Onceasa | 43.7 | 15.7 | 3.3 | -20.9 | 3.0 | M |
| USR95 | Onceasa | 37.2 | 13.7 | 3.2 | -21.5 | 4.5 | M |
| USR96 | Onceasa | 40.2 | 14.7 | 3.2 | -21.5 | 5.2 | M |
| USR97 | Onceasa | 40.9 | 14.7 | 3.3 | -21.8 | 4.8 | F |
| USR98 | Onceasa | 41.3 | 14.9 | 3.2 | -22.1 | 4.9 | M |
| USR99 | Onceasa | 39.1 | 13.9 | 3.3 | -22.5 | 6.1 | F |
| USR100 | Onceasa | 42.5 | 15.3 | 3.2 | -21.1 | 3.6 | M |
| USR101 | Ferice | - | - | - | - | - | |
| USR102 | Ferice | - | - | - | - | - | |
| USR103 | Ferice | - | - | - | - | - | |
| USR104 | Ferice | - | - | - | - | - | |

| | | | | | | | |
|---|---|---|---|---|---|---|---|
| USR105 | Ferice | - | - | - | - | - | |
| USR106 | Ferice | - | - | - | - | - | |
| USR107 | Ferice | - | - | - | - | - | |
| USR108 | Lelici | 40.3 | 14.1 | 3.3 | -22.0 | 6.4 | M |
| USR109 | Lelici | 40.5 | 14.3 | 3.3 | -21.1 | 4.0 | F# |
| USR110 | Lelici | 39.0 | 14.0 | 3.3 | -20.6 | 2.7 | F |
| USR111 | Lelici | 42.4 | 15.0 | 3.3 | -21.0 | 4.0 | M# |

**Table S2**. Details of $^{14}C$ dates and molecular estimated ages of samples for which nuclear genome and mitogenome data have been generated. $^{14}C$ dates for USR11, USR22, USR25, USR65 are from Naito et al. (2020).

| **$^{14}C$ dates and molecular estimated ages of samples** | | | | |
|---|---|---|---|---|
| **Sample ID** | **Sampling cave** | **Uncalibrated $^{14}C$ date/Molecular tip date*** **(yrs BP)** | **Calibrated dates (95.4%) (cal BP) / 95% credibility interval for molecular estimates*** | **$^{14}C$ lab number** |
| USR10 | Măgura | 29,903 * | 26,830 - 32,217* | |
| USR11 | Măgura | 24,615 ± 101 | 28,680 - 29,120 | ETH-87394 |
| USR22 | Cioclovina | 40,567 ± 594 | 42,855 - 44,450 | ETH-87395 |
| USR23 | Cioclovina | 32, 882 ± 233 | 36,594 - 38,374 | ETH-87716 |
| USR25 | Cioclovina | 43,119 ± 807 | 44,474 - 47,414 | ETH-87396 |
| USR65 | Răsuflătoarei | 31,005 ± 183 | 34,816 - 35,913 | ETH-95736 |
| USR67 | Răsuflătoarei | 35,751 * | 32,078 - 38,945 * | |
| USR91 | Onceasa | 31,191 ± 200 | 35,195 - 36,119 | ETH-87392 |
| USR92 | Onceasa | 35,041 * | 31,429 - 38,129 * | |
| USR109 | Lelici | 26,358 ± 119 | 30,295 - 30,955 | ETH-87393 |
| USR111 | Lelici | 29,792* | 26,993 - 32,142 * | |

**Table S3**. Best partitioning scheme selected by PartitionFinder for Bayesian mitochondrial tip dating analysis.

| Partition | Best Model | Mitochondrial DNA elements |
|---|---|---|
| 1 | HKY+X | Lys, Ile, His, Asp, Arg, Gly, Asn, 16s_pt2, Ser1, Glu, Phe, Trp, Ser2, Leu2, 12s, 16s_p1, ND6_CP2, ND2_CP1, ATP8_ATP6_CP3, ND5_CP1 |
| 2 | HKY+G+X | Ala, Gln, Pro, ND6_CP3, ND3_CP3, ND4L_CP3, COX3_CP3, ND4_CP1, COX2_CP3, Tyr, Leu1, Thr, D-loop, COX1_CP3, ATP8_ATP6_CP2 |
| 3 | K80 | ND3_CP1, Cys, COX3_CP1, Val, CYTB_CP1, COX1_CP1, ND1_CP1, Met, COX2_CP1, ND4L_CP1 |
| 4 | HKY+X | ND4_CP2, COX1_CP2, COX3_CP2, CYTB_CP2, ND4L_CP2, ND1_CP2, ND2_CP2, COX2_CP2, ND5_CP2, ND3_CP2, ATP8_ATP6_CP1 |
| 5 | HKY+G+X | ND6_CP1, ND5_CP3, ND2_CP3, CYTB_CP3, ND1_CP3, ND4_CP3 |

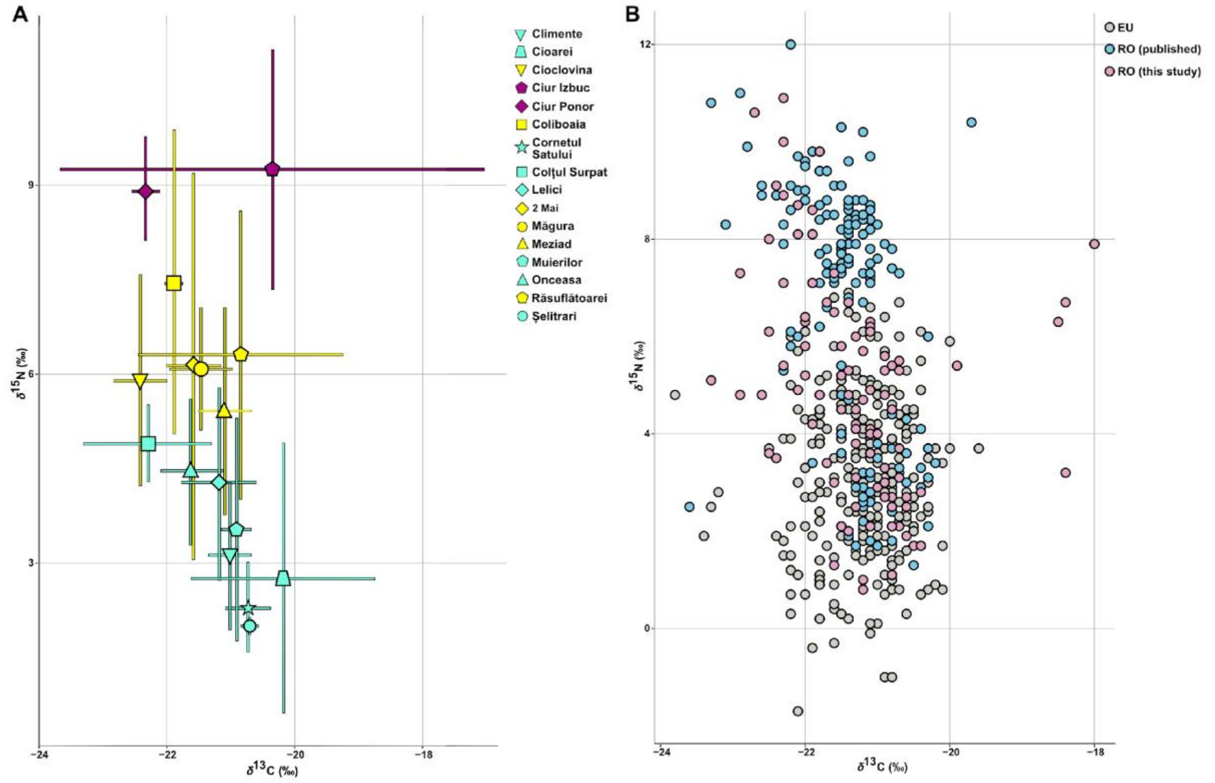


**Figure S1. Comparative isotopic patterns of Romanian cave bears and European sites. A.** Plot depicting isotopic patterns among cave bears from 16 sites across Romania. Igrița and Stogu caves are not represented due to insufficient sample sizes (n = 1). The highest variability in both $\delta^{13}$C and $\delta^{15}$N values is recorded in Ciur Izbuc (NW RO, Apuseni Mts, intra-Carpathian basin), followed by 2 caves, Răsuflătoarei and 2 Mai caves located in SW RO, Banat Mts (intra-Carpathian basin). The lowest variability is depicted in caves located: on the outer side of the Romanian Carpathians (i.e. Cornetul Satului, Șelitrari – likely due to small sample size n=2; but also in better sampled caves like Muierilor and Cioarei); in the intra-Carpathian basin at high altitudes > 1,000 m a.s.l. (see also Figure 1). Colours correspond to the three adult diet types as shown in Figure 1. **B.** Scatter-plot of bulk collagen $\delta^{13}$C and $\delta^{15}$N values of published individual cave bears from Europe (283 individuals from Austria, Belgium, France, Germany, Italy, Poland, Slovakia, Spain, Switzerland, dataset from Bocherens, 2019), from published Romanian sites (155 individuals, dataset from Robu et al. 2013; 2017) and from this study (95 individuals).

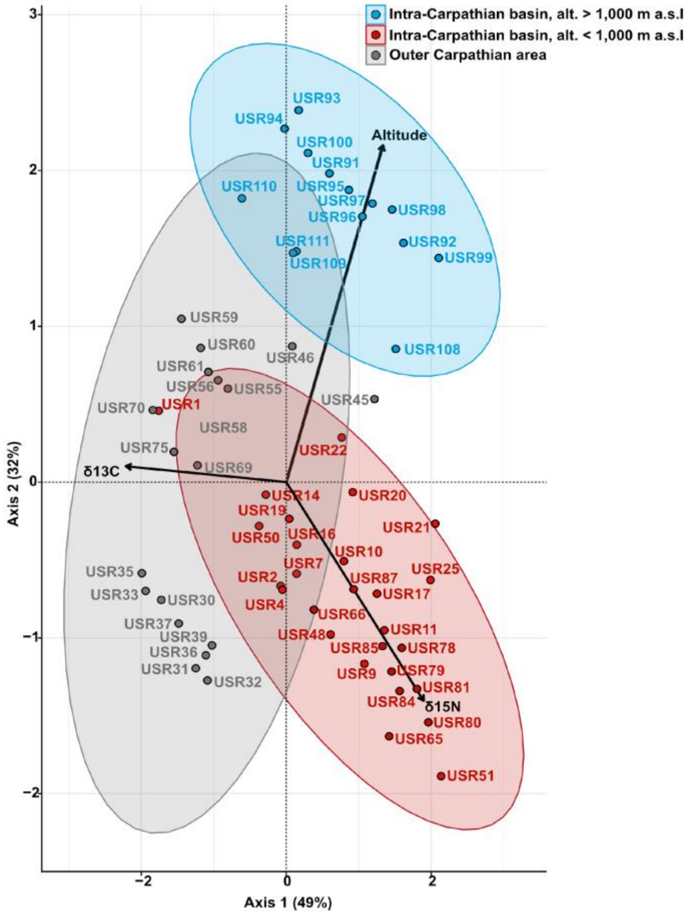


**Figure S2. Figure 3. PCA of cave bear isotopic values and altitude in Romanian Carpathian caves.** PCA based on bulk collagen $\delta^{13}$C and $\delta^{15}$N values and altitude for cave bears from 14 caves of the Romanian Carpathians (n=89). USR: sample code. Vectors correspond to explanatory variables. For the sake of clarity, only samples that were projected better than 0.75 cos$^2$ on the 2 axes are listed.

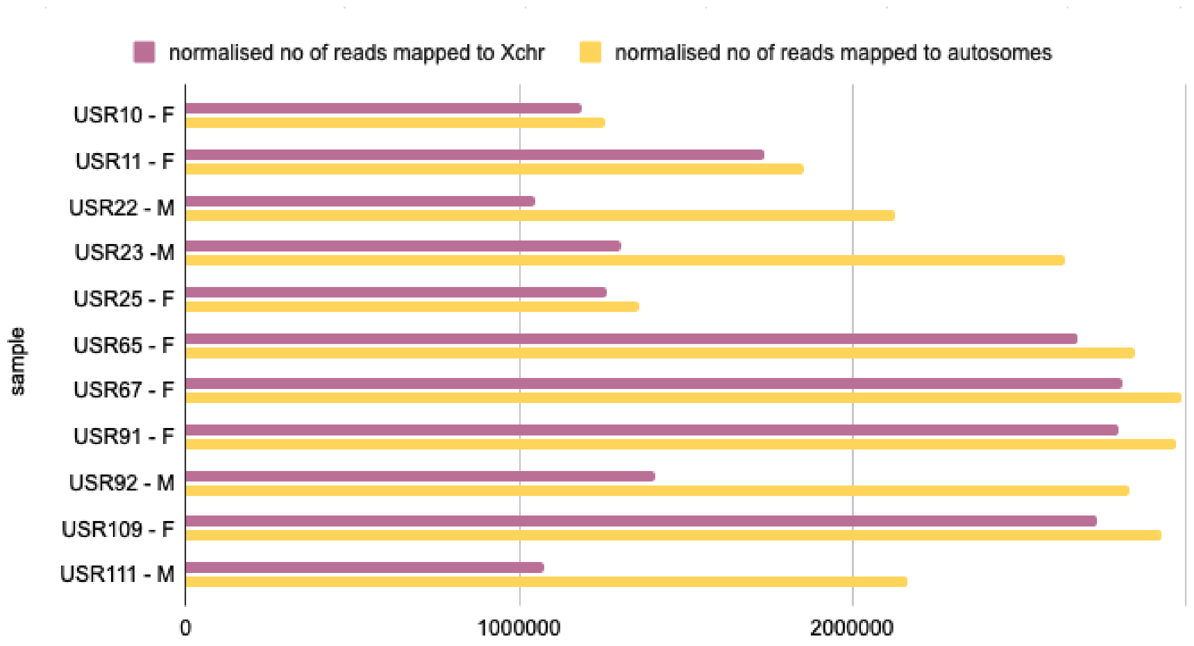


**Figure S3.** Normalized number of reads mapped to the X chromosome versus autosomes. Individuals were classified as females when the ratio of mapped reads to X chromosomes against autosomes was approximately equal, and as males when the ratio was approximately 0.5.
